# Substrate-dependent epistasis probes active site intramolecular wiring

**DOI:** 10.64898/2026.07.02.736193

**Authors:** Karol Buda, Charlotte M. Miton, Clara Vogt, Nobuhiko Tokuriki

## Abstract

Enzyme adaptation toward novel substrates involves the rewiring of intramolecular residue networks, yet how this rewiring differs across multiple substrates, and how it underpins functional trade-offs and promiscuity, remains poorly understood. Here, we profile all 64 combinations of six key mutations in a phosphotriesterase across nine structurally diverse substrates spanning three chemical classes (organophosphates, esters, and lactones), thus generating a multi-dimensional map of epistasis and promiscuity within the phosphotriesterase’s active site. We developed a statistically robust reference-based analysis pipeline incorporating error propagation and significance testing to move beyond global epistatic trends and resolve idiosyncratic, substrate-dependent intramolecular wiring in specific genetic backgrounds. Simulations confirm that this pipeline reliably identifies genuine higher-order epistatic interactions while minimizing false positives. We reveal that intramolecular network wiring varies substantially between substrates – even within the same chemical class – with notable divergences between the adaptive target substrate 2-naphthyl hexanoate and its shorter-chain ester analogs. Key higher-order networks, including *d233***E**/*h254***R**/*l271***F** and *l271***F**/*f306***I**/*i313***F**, exhibit substrate-specific epistatic signatures that discriminate between subtle structural features such as acyl chain length, leaving group identity, and heteroatom substitution. These substrate-dependent rewiring events account for observed functional trade-offs, particularly the strong anti-correlation between the adaptive and native substrates. Collectively, these findings demonstrate that comprehensive cross-substrate epistatic profiling, paired with rigorous statistical analysis, provides a powerful framework for dissecting the molecular basis of enzyme promiscuity and the trade-offs that define adaptive evolution.

**Significance of the study:** Enzymes are biological catalysts whose function depends on precisely coordinated networks of interacting amino acids. Understanding how these networks change as enzymes evolve to recognize new molecules is fundamental to understanding and predicting evolution. Here, we developed a statistical framework to map these networks across multiple molecules simultaneously, revealing how subtle differences in molecular shape drive functional trade-offs during evolution. These insights could inform the rational design of enzymes with tailored specificities for industrial and therapeutic applications.

## Introduction

Enzymes are far more than the sum of their individual residues; they consist of finely-tuned intramolecular networks where amino acids are “wired” together to coordinate complex catalytic functions (Miton et al., 2021). Adaptive mutations that alter enzyme function towards novel substrates do so by modulating these intramolecular networks. Furthermore, enzyme evolution toward new substrates is often accompanied by shifts in catalytic efficiency across various other substrates. Activity toward the original (native) substrate is often reduced, *i.e.*, results in a functional trade-off, while activity to other promiscuous substrates may also change, leading to coordinated co-evolution of substrate specificity – all underpinned by intricate changes in intramolecular network wiring. The interplay between network rewiring, adaptation, and function across multiple substrates represents a key mechanism that drives enzyme evolution. Ideally, such multi-substrate network rewiring would be characterized through structural biology approaches, however, obtaining high-resolution structures for all relevant variants in an evolutionary trajectory remains experimentally challenging. In addition, one would also require an understanding of how these enzyme variants interact with all relevant substrates and, more importantly, with their transition states, which presents a significant experimental barrier. Many mutational effects may also be invisible in static crystal structures, as their impacts can be dynamic and involve subtle changes in the intra- and inter-molecular interactions that are not readily captured *in crystallo* (Naganathan, 2019; Summers et al., 2023). Thus, it is necessary to explore alternative approaches in order to capture these rewiring events across several substrates.

Combinatorial mutational analysis has become a powerful approach to illuminate how the wiring of intramolecular networks helps facilitate enzyme functions. By identifying key mutations in an adaptive process, one can conduct rigorous functional interrogation of variants that comprehensively sample a sub-space of mutational combinations, enabling the quantification of mutational epistasis, including higher-order interactions involving three or more mutations (Weinreich et al., 2006). Contemporary analyses of such landscapes utilize reference-free analysis (RFA), which captures the overall architecture of the entire genotype-phenotype map. RFA is particularly powerful because it mitigates the dangers of attributing epistasis to experimental uncertainty by capturing average epistatic trends across the landscape (Park et al., 2024; Poelwijk et al., 2016). Using variations of RFA, previous studies have hinted at how intramolecular networks may be formed and broken during evolution towards a specific substrate, and how epistatic interactions differ across other substrates or cofactors (Anderson et al., 2021; Lozovsky et al., 2020; Mira et al., 2015; Yang et al., 2019; Zhang et al., 2012).

The main drawback, however, of using these averaged epistasis terms obtained through RFA, is that they obscure the idiosyncratic wiring of residues in specific genotypes; the magnitude of a specific interaction is captured across all representative genotypes. Thus, RFA-based epistatic effects cannot be attributed to specific interactions that define the ligand-network wiring in a given genotype, limiting our ability to explore intramolecular wiring. A solution is offered by reference-based analysis (RBA), which has the potential to decode the wiring of idiosyncratic genotypes (Buda et al., 2023). Indeed, RBA coupled with structural analyses has enabled understanding of pairwise network formation during adaptive evolution for several enzymes (Campbell et al., 2016; Dellus-Gur et al., 2015; González et al., 2016; Judge et al., 2024; Sunden et al., 2015). However, RBA has historically been hindered by the intrinsic weaknesses of relying on a specific reference state (typically the wild-type) and, thus, RBA is highly sensitive to the propagation of experimental error in the calculation of epistasis at higher-orders. Indeed, recent empirical work by Camacho-Mateu *et al*. demonstrated the difficulty in detecting higher-order epistatic effects in interactions between microbes (Camacho-Mateu et al., 2026). Thus, the development of an RBA pipeline that can assess the significance of epistatic terms, particularly at higher-orders, for enzyme function across multiple substrates, is essential to define the complexity of intramolecular networks and their differential wiring in the context of enzyme promiscuity.

We hypothesize that studying epistatic network variability across different substrates can provide molecular insights, particularly the rewiring of intramolecular networks during mutational accumulation, into functional trade-offs and co-evolution towards multiple substrates. In this study, we demonstrate how epistatic networks can serve as a foundation for structural speculations that elucidate the substrate-dependent wiring of a model enzyme: phosphotriesterase (PTE). Previously, we evolved PTE from its native substrate ethyl-paraoxon (EPO), an OP pesticide xenobiotic, towards hydrolysis of the ester 2-naphthyl hexanoate (2NH) over 18 rounds of evolution (Kaltenbach et al., 2016; Tokuriki et al., 2012; Wyganowski et al., 2013). From the 18 accumulated mutations, additional research deeply characterized the mechanism by which six key mutations help catalyze 2NH (Campbell et al., 2016; Miton et al., 2020). Here, the activity of all 64 combinations of these six mutations was measured across nine substrates belonging to three distinct chemical classes, providing a multi-dimensional map underpinning the interplay between epistasis and promiscuity within the PTE active site. We developed a statistically robust RBA pipeline that includes error propagation calculations to filter for significant epistatic effects, allowing us to move beyond global trends of RFA. These results reveal how combinatorial landscapes, when profiled across diverse substrates, can uncover subtle but critical differences in active site intramolecular wiring, providing a structural rationale for the evolution of promiscuity and the trade-offs that define enzyme adaptation.

## Results

### Experimental methods and analytical pipeline

Using site-directed mutagenesis, we previously constructed 64 mutants consisting of wt (high EPO activity) and derived (high 2NH activity) states across six positions: *d233***E**, *h254***R**, *l271***F**, *l272***M**, *f306***I**, *i313***F** (**Fig. 1a)** (Miton et al., 2020); *italic* font represents ancestral states while **bold** font represents derived states. We have previously measured the activity across the entire combinatorial landscape for *p*-nitrophenyl butyrate (Miton et al., 2020), as well as for 2NH (Buda et al., 2023). In this study, we expanded our substrate set to a total of nine substrates: three esters [2NH, *p*-nitrophenyl butyrate, and *p*-nitrophenyl acetate], four OPs [EPO, methyl-paraoxon (MPO), ethyl-parathion (MPT), and methyl-parathion (MPT)], and two lactones [dihyrocoumarin (DHC), and thiobutyl-butyrolactone (TBBL)] (**Fig. 1b**). OPs represent the native substrate class for PTE, esters were the targeted class of substrates during directed evolution, and lactones are the putative ancestral substrate class for PTE (Afriat-Jurnou et al., 2012). Several substrates in each class were chosen to investigate how changes to the intramolecular network *via* mutations may reveal subtle differences between functional changes for substrates belonging to the same class, *i.e.*, substrate promiscuity, in addition to different substrate classes, *i.e.*, catalytic promiscuity. Activity of all substrates was measured *via* colorimetric detection of hydrolyzed products upon addition of substrate to *Escherichia coli* lysates containing expressed PTE variants (see **Methods**).

**Figure 1.**
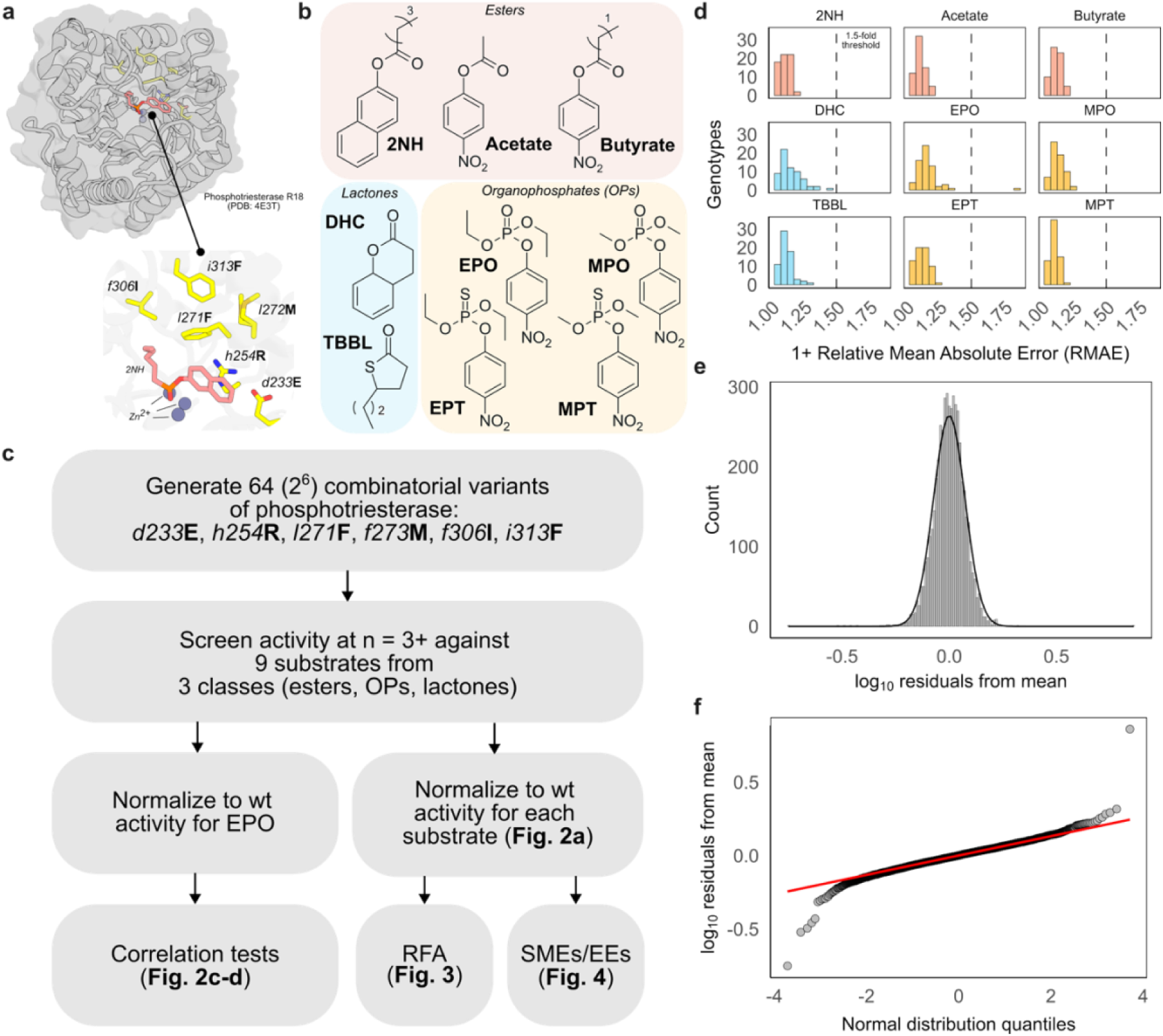
Overview of the model system, data processing pipeline, and data quality. a, The crystal structure of round 18 phosphotriesterase (PTE-R18; PDB: 4E3T) complexed with hexyl(naphthalen-2-yloxy)phosphinic acid, a 2-naphthyl hexanoate (2NH) transition state analogue, highlighted in salmon, and six yellow residues constituting the explored mutations in the study. **b,** Chemical structures of all substrates used in the study: esters [2NH, p-nitrophenyl butyrate, and p-nitrophenyl acetate], four OPs [ethyl-paraoxon (EPO), methyl-paraoxon (MPO), ethyl-parathion (MPT), and methyl-parathion (MPT)], and two lactones [dihyrocoumarin (DHC), and thiobutyl-butyrolactone (TBBL)]. **c,** Data processing pipeline of combinatorial data. **d,** Error ranges for the log_10_ mean activity of each mutant for every substrate, with a 1.5-fold threshold denoted. **e,** Visualization of the log-normal distribution of experimental error for all replicates. **f,** Quantile-quantile plot demonstration of log-normality with minimal outliers.

To investigate the robustness of the *in vitro* lysate assays for the computation of epistasis, we first assessed the measurement errors across substrate classes. We collected 4872 measurements, consisting of biological (n = 3–10) and technical (n = 3) replicates for 576 conditions (64 genotypes across nine substrates). Replicate measurements were averaged for each genotype-substrate combination, log_10_ transformed, and normalized to a reference wt activity (**Fig. 1c** and **Methods**). Measurements for each genotype-phenotype pair across all substrates exhibited coefficients of variation (CVs) ranging from 5% to 142% (mean = 16%; median = 15%). To compare with our previously used 1.5-fold error threshold of significance (Buda et al., 2023), we computed a relative mean absolute error (RMAE; see **Methods**). RMAEs ranged from 1.03–1.83-fold (mean = 1.13-fold; median = 1.12-fold), much lower than the previously utilized 1.5-fold threshold, and were similarly distributed across and within substrate classes (**Fig. 1d**). We found that replicate measurements across all substrates were log-normally distributed around each genotype’s mean activity (mean = 0.00, range = [-0.75, 0.86], standard deviation = 0.07) (**Fig. 1e-f**). Thus, we found the data suitable for downstream assessment of cross-substrate correlations and the extraction of epistatic terms.

### Combinatorial landscape phenotypes differ across all substrates

First, we aimed to explore the similarities and differences in effects that all mutations, across all genetic backgrounds, played across all measured substrates. To do so, we began by investigating the functional contribution of mutations across all substrate-dependent combinatorial landscapes. We separated activities by substrate, and then normalized the activities to the wt background value, resulting in log_10_ fold-changes relative to wt for each respective substrate (**Fig. 1b**). As expected, mutational accumulation during directed evolution resulted in an increased 2NH activity (the target) and other ester substrates, accompanied by a decrease in EPO and other OP activity (the native function) (**Fig. 2a**). Interestingly, we observed that the activity changes for the two lactones differed from one another, with an overall decrease in activity for DHC, while TBBL showed some increase in activity, particularly for genotypes containing *h254***R** (**Fig. 2a**).

**Figure 2.**
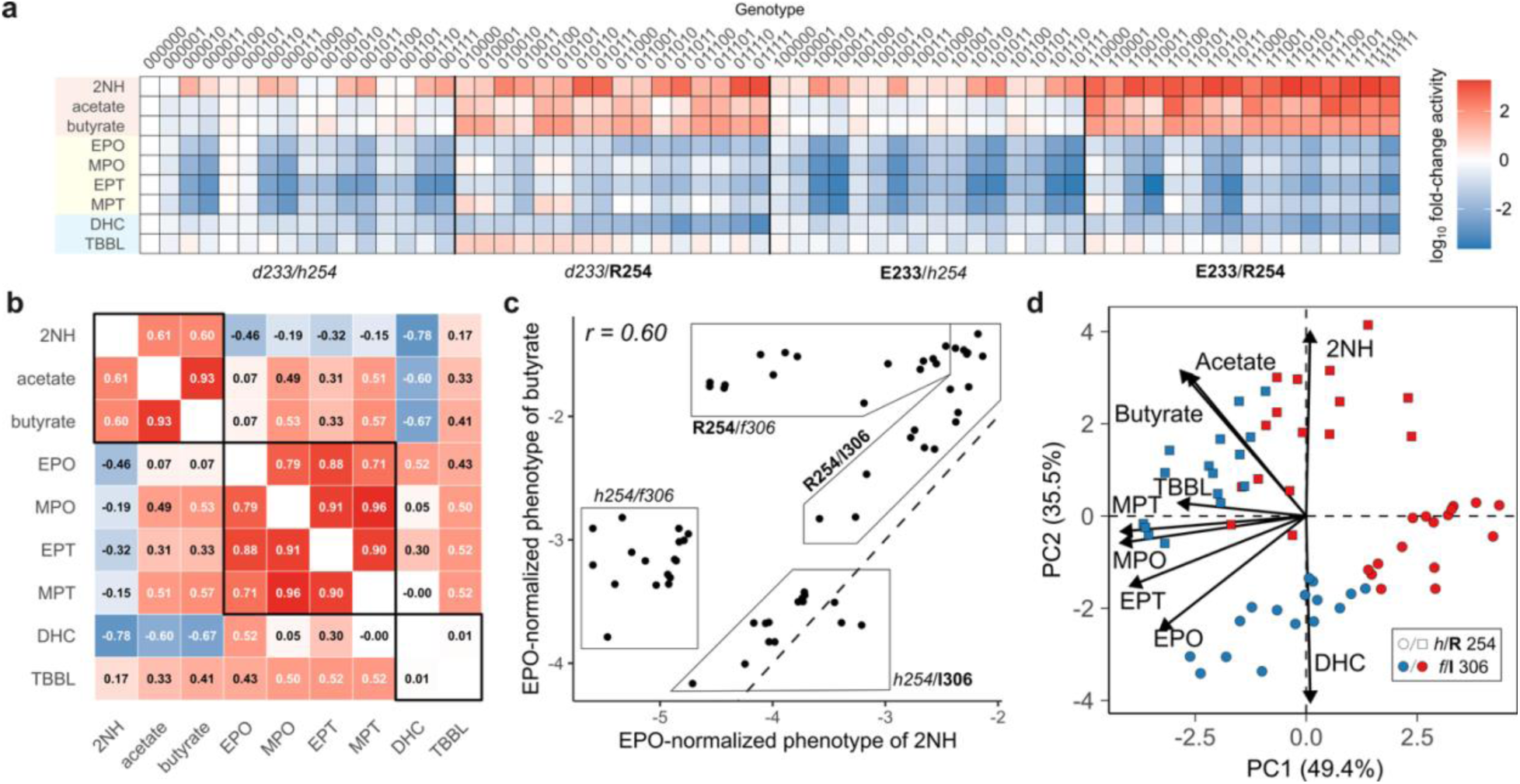
**Mutations differentially affect cross-substrate activity in the phosphotriesterase combinatorial landscape**. **a,** log_10_ fold-change in activity for each substrate across 64 combinatorial mutants, ordered by presence of the 233/254 pair. **b,** Pearson correlation coefficient matrix between log_10_ activities normalized to wt ethyl-paraoxon (EPO) function across substrates: three esters [2-naphthyl hexanoate (2NH), p-nitrophenyl butyrate, and p-nitrophenyl acetate], four organophosphates [ethyl-paraoxon (EPO), methyl-paraoxon (MPO), ethyl-parathion (MPT), and methyl-parathion (MPT)], and two lactones [dihyrocoumarin (DHC), and thiobutyl-butyrolactone (TBBL)]. **c,** Correlation between 2NH and butyrate activity highlights 2NH-specific contribution of the mutation at residue 306. **d,** Principal component analysis of activity-substrate data with identity of *h254***R** (circles *versus* squares) and *f306***I** (blue *versus* red) highlighted.

We investigated these effects further by normalizing activities for all substrates to the wt activity for EPO, then compared the Pearson correlation coefficients (*r*) of activity changes across the entire combinatorial landscape between each substrate pair (**Fig. 2b**). For within-substrate-class correlations, we found that OPs were positively correlated with *r* = [0.71, 0.96], and showed the greatest discrimination between ethyl and methyl substituted OPs. Notably, EPO had the lowest maximum correlation to all OPs relative to any other OP, suggesting a specific disruption in activity for the native EPO substrate over other OPs. The two *p*-nitrophenyl substituted esters showed a very strong correlation of 0.96; however, both shorter chain esters showed weaker correlations with 2NH at Pearson values of 0.61 and 0.60 for acetate and butyrate, respectively. This weaker correlation appeared to be mostly affected by the contribution of *f306***I** (**Fig. 2c**). It is apparent that *h254***R** is beneficial for all ester hydrolysis, and the most important factor dictating functional improvement in butyrate, while the presence of either *h254***R** or *f306***I,** or their combination, appears to specifically improve 2NH activity (**Fig. 2c**). This suggests that distinct enzyme-substrate interactions are formed with (*i*) the longer hexanoate tail or (*ii*) the naphthyl group, both absent from the acetate and butyrate. Finally, we found no correlation between DHC and TBBL (*r* = 0.01), indicating substantially different mechanisms by which this six-residue network supports the hydrolysis of lactones.

Comparisons of between-substrate-class correlations revealed weak anticorrelations (*r* = [−0.15, −0.46]) between 2NH and all OPs, as expected, since this constitutes the 2NH/POE trade-off previously observed (Tokuriki et al., 2012). Interestingly, we observed a positive correlation between methyl OPs and the *p*-nitrophenyl esters, a weaker positive correlation with EPT, and no correlation with EPO (**Fig. 2b**). This may suggest that the means by which the six residues help facilitate the hydrolysis of *p*-nitrophenyl containing substrates is similar, particularly for the smaller methyl OPs which may be positioned in the active site similarly to acetate and butyrate. DHC revealed strong anticorrelations with all esters (*r* = [−0.60, −0.78]), no correlation with methyl OPs, and a positive correlation with ethyl OPs (**Fig. 2b**). Finally, TBBL revealed a preferential correlation for butyrate, with no apparent correlations to any other substrate (**Fig. 2b**). For both lactones, the exact role of these six mutations in facilitating hydrolysis is unknown, thus it is difficult to speculate on these trends. Nonetheless, probing all genotype-phenotype pairs with these nine substrates highlights potential differences in the mechanisms of hydrolysis within and across substrate classes.

To capture the multi-dimensional nature of the multi-substrate combinatorial landscapes, and address the functional contributions of all mutations, especially by 254 and 306, we resorted to a principal component analysis (PCA) on all activity data. We found that principal components 1 (PC1) and 2 (PC2) jointly accounted for 84.9% of the variance. PC1 strongly distinguishes the loading vectors of 2NH and DHC from other substrates, while PC2 separates 2NH and DHC into opposite “poles,” while also separating the esters from the OPs and TBBL (**Fig. 2d**). Next, we investigated how genotypes are partitioned across the PCs, and found that PC1 appears to separate genotypes by their *f306***I** identity, with 2NH and DHC being more tolerant of, or preferring **I***306*, while PC2 partitioned genotypes by their *h254***R** identity, demonstrating preference for **R***254* in the esters, TBBL, methyl-substituted OPs, EPT, EPO, and DHC, in that order (**Fig. 2d**). We found that the cluster of genotypes containing *h254* and **I***306* was not associated with any loading vectors, indicating that **I***306* relies on other mutations for large functional improvement across all substrates, including 2NH. The *p*-nitrophenyl ester loading vectors were strongly associated with **R***254* and *f306* genotypes, while the 2NH loading vector was oriented towards the **R***254* and **I***306* cluster. Notably, the directionality and magnitude of the shorter chain esters was nearly identical, suggesting that the intramolecular network supporting their hydrolysis is highly similar, also revealed by their positive correlation for activity (**Fig. 2b**). Thus, PCA revealed that *f306***I** in isolation is a good predictor of poor activity across most substrates, while *h254***R** three main clusters − DHC, the esters, and OPs with TBBL, and finally *h254***R/***f306***I** − is a good predictor of high 2NH activity, as well as low DHC activity. Other mutations were broadly distributed across the PCs and did not reveal additional structure in the data.

### Threshold epistasis and reference-free analysis

To capture the underlying functional variance across the six-residue network, we first employed RFA. We used RFA to (*i*) capture the average effect of a mutation or epistatic interaction across all genotypes and (*ii*) assess the overall epistatic complexity of the entire landscape by identifying the minimum number of mutational terms, regardless of a specific genetic starting point, required to reconstruct the observed phenotypes. Before extracting these terms, we addressed the potential for threshold epistasis – the phenomenon where the functional contributions of mutations on an underlying linear trait are distorted by a non-linear filter, such as enzyme stability or experimental detection limits. While we observed significant improvements in model fit for all substrates upon applying a sigmoidal transformation (p = 0.015; df = 26), the mean increase in the out-of-sample *R^2^* was negligible (Δ*R^2^* = 0.007; **Fig. S1**). Furthermore, the coefficients of the sigmoidal model that resulted in the best fits varied across substrates. Given that non-specific epistasis is typically assumed to reflect substrate-independent biological constraints, like protein stability, the need for substrate-specific transformations indicated that these effects are better explained by specific mutational interactions rather than a global threshold model (Dupic et al., 2024). Consequently, we proceeded with an RFA of the untransformed data.

We quantified the complexity of each landscape using the T_90_ metric, representing the minimum number of reference-free terms needed to achieve an *R^2^* of 90% in model predictions. Across all nine substrates, we found that the functional variance was sufficiently captured by a 2^nd^ order model, with a mean T_90_ = 14.6 terms and a median T_90_ = 15 terms (**Fig. 3**). 1^st^ order terms accounted for the majority of the variance for every substrate, and second-order terms were only incorporated after these primary effects were exhausted. Intriguingly, for all substrates except TBBL, a brief decrease in the out-of-sample *R^2^* was observed when certain new terms were incorporated, before the accuracy eventually recovered. This suggests that idiosyncratic genotypes within the landscape can temporarily distort global predictions of partial models, indicating that individual “average” epistatic effects are sensitive to the unique intramolecular networks formed by specific mutations in certain backgrounds. Nonetheless, a comprehensive understanding of all landscapes was achieved with 2^nd^ order terms; this does not exclude the presence of higher-order epistasis, but rather demonstrates that the majority of the variance can, on average, be explained by single and pairwise mutations.

**Figure 3.**
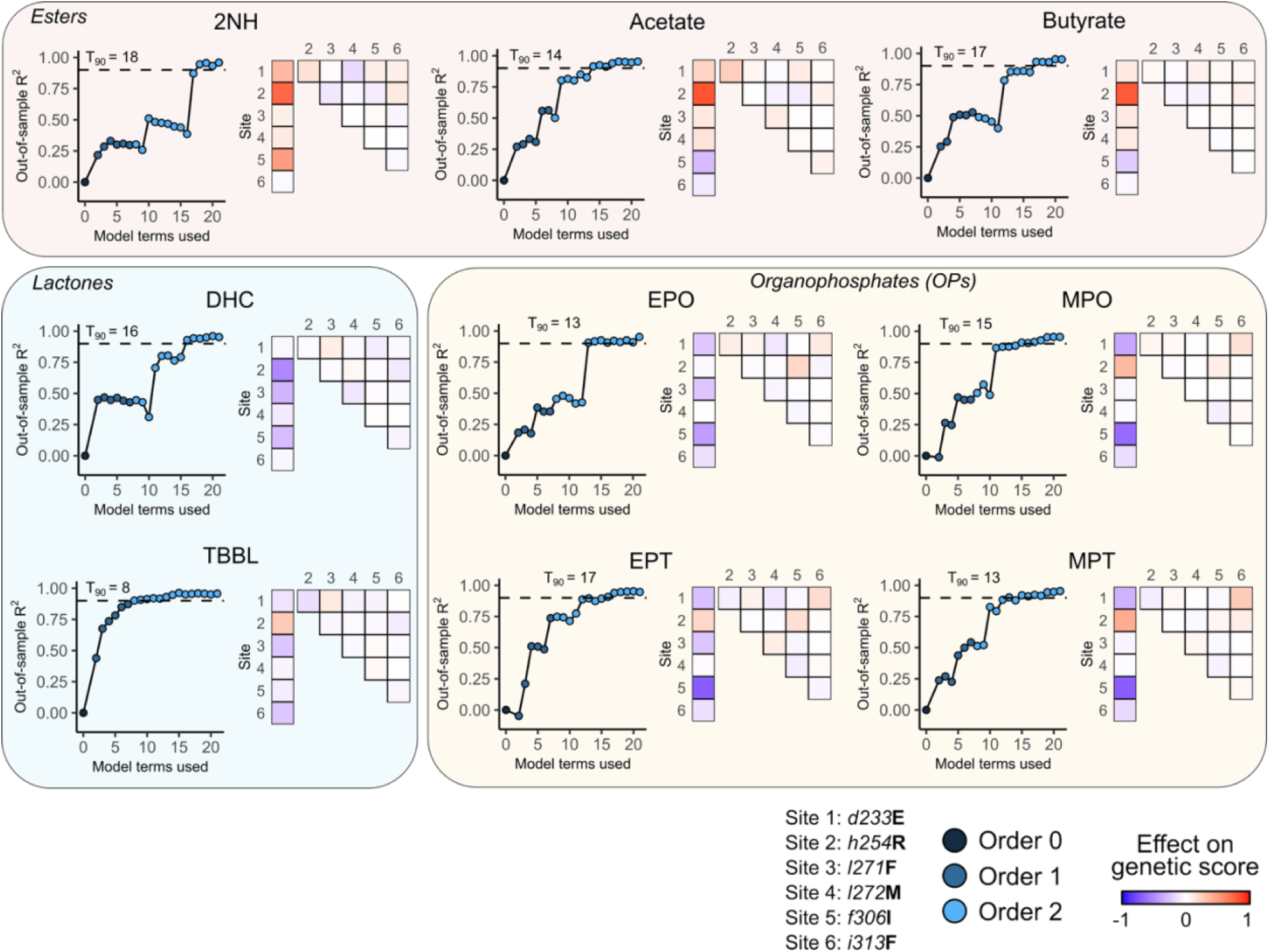
Reference-free analysis reveals key differences and similarities in epistasis across nine substrates. Analyses were performed on substrates from three classes: three esters [2-naphthyl hexanoate (2NH), p-nitrophenyl butyrate, and p-nitrophenyl acetate], four organophosphates [ethyl-paraoxon (EPO), methyl-paraoxon (MPO), ethyl-parathion (MPT), and methyl-parathion (MPT)], and two lactones [dihyrocoumarin (DHC), and thiobutyl-butyrolactone (TBBL)]. Left panels: analyses of minimum model terms required for 90% R^2^ on non-training set data, with model order of the next parameter annotated in shades of blue. Right panels: heatmaps of the effect of mutations at each site on the genetic score; blue are negative effects and red are positive.

Moreover, the RFA confirmed broad functional trends from the previous analyses: the *f306***I** mutation was found to contribute positively almost exclusively to 2NH activity, while the *h254***R** mutation showed a positive contribution across all substrates except for the native substrate EPO, as well as DHC (**Fig. 3**). RFA also helped reveal more subtle effects: we found that *l271***F** appeared detrimental specifically in the context of ethyl-substituted OPs, that *l272***M** showed slight ester preference with minimal trade-off, and that the average first-order effect of *i313***F** was negative or neutral across all substrates, but exhibited positive pairwise effects with some mutations for esters and OPs.

### Reference-based analysis with error propagation identifies substrate-dependent epistatic effects

In order to connect epistatic terms to specific interactions between amino acids in the enzyme structure, we first needed to compute these terms in specific genetic contexts. To do so, we turned to RBA. In contrast to RFA, RBA computes the functional contribution of mutations and epistasis in specific genetic backgrounds, rather than across all genotypes across the landscape (see **Methods**). Thus, RBA is necessary to (*i*) interrogate the presence of epistatic wiring, including higher-order epistasis, in idiosyncratic genotypes, *i.e.*, outliers, where wiring is unique to that genotype relative to others in the landscape and (*ii*) to structurally map, compare, and contrast the idiosyncratic epistatic interactions for each substrate within a specific genotype. For each substrate, we first computed the single mutational effect (SME) for each of the six positions across all backgrounds (see **Methods**). We have previously demonstrated how the spread in SMEs and difference between the wt SME and the average SME across all genotypes are indicators of idiosyncratic effects across the landscape (Buda et al., 2023). As expected, mirroring the RFA analysis, *f272***M** and *i313***F** showed relatively minor spread and difference between wt and average effect (**Fig. 4a**). We found that *d233***E** showed a greater spread, with positive effects in esters and negative effects in OPs, though still exhibited similar effects in the wt as it did on average. We observed the strongest idiosyncrasy in *h254***R**, *l271***F**, and *f306***I**. Interestingly, although all *h254***R** SMEs in esters were positive, we found that the effect was initially weaker in the wt than on average for 2NH and acetate, but not butyrate, indicating epistatic network formation in the former two substrates, but not the latter. The effect of *h254***R** was negative for ethyl-substituted OPs in the wt, but positive in methyl-substituted OPs, and was on average stabilized by other mutations for all substrates (**Fig. 4a**). For *l271***F**, the SMEs were markedly more negative in the wt for thiol-substituted OPs than on average, while other substrates showed relatively similar effects (**Fig. 4a**). Finally, *f306***I** showed a strong positive effect for 2NH in the wt, though it became weaker on average, something we observed previously in the context of higher-order network formation (Buda et al., 2023). In contrast, *f306***I** was strongly negative for all OPs in the wt background, but became more tolerant on average with the accumulation of other mutations (**Fig. 4a**). The idiosyncratic effects across these positions demonstrate the need to capture and understand the wiring that modulates their effects across this panel of substrates.

**Figure 4.**
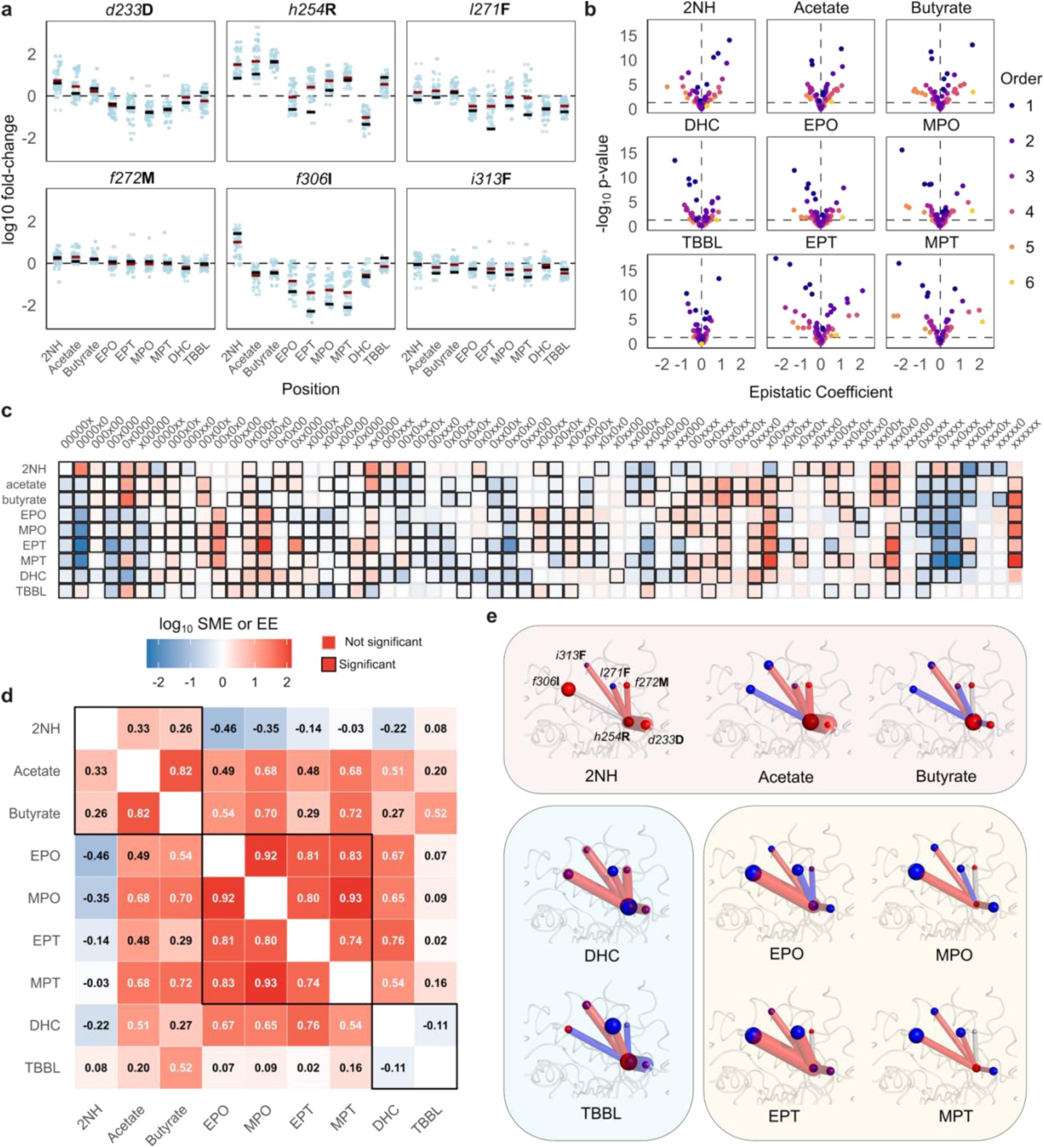
Reference-based analysis with error propagation reveals significant, higher-order epistatic effects. a, The log_10_ fold-change in activity of each single mutational effect (SME) of the six mutations across each of the 32 genetic backgrounds, partitioned by substrates: three esters [2-naphthyl hexanoate (2NH), p-nitrophenyl butyrate, and p-nitrophenyl acetate], four organophosphates [ethyl-paraoxon (EPO), methyl-paraoxon (MPO), ethyl-parathion (MPT), and methyl-parathion (MPT)], and two lactones [dihyrocoumarin (DHC), and thiobutyl-butyrolactone (TBBL)]. Mean effect of the mutation is represented by the dark red bar and SME in the wt-background is represented by the black bar. **b,** Volcano plots denoting significant SMEs and epistatic effects (EEs) based on p < 0.05, with sign and magnitude also shown. **c,** A heat map of the sign and magnitude of all log_10_ SMEs and EEs for each substrate, with transparent borders indicating not significant coefficients and opaque borders indicating significant ones. **d,** SMEs and pairwise EEs represented in the phosphotriesterase active site, with spheres representing SMEs and lines representing EEs. Thickness corresponds to magnitude and color to sign, with positive interactions shown in red, negative in blue, and not significant in grey. **e,** Pearson correlation coefficient matrix of all SMEs and EEs between substrates.

To enable assessment of the wiring between the six positions in different substrate contexts, we computed the wt SME and epistatic effects (EEs) for all mutational combinations; wt EEs represent functional effects of an interaction between 2-6 residues (see **Methods**). Given the abundance of replicates for lysate data across all substrates, when computing SMEs and EEs, we used the extracted standard deviation values for the mean activity of each genotype-substrate pair for error propagation analysis. To evaluate the significance of each SME and EE, we also calculated effective sample sizes for each EE by computing the harmonic mean of all replicates for each constituent measurement of the SME and EE, *e.g.*, the *d233***E**/*h254***R** EE for 2NH would have an effective sample size based on the harmonic mean of wt PTE, *d233***E**, *h254***R**, and *d233***E**/*h254***R** replicate measurements (see **Methods**). Using propagated standard deviation and effective sample sizes, we computed t-values and p-values for each SME and EE, and assessed their significance based on a threshold of p < 0.05 (**Fig. 4b**). We found that 63.5% (360/567) of the computed SMEs and EEs across the nine substrates were deemed significant. We observed significant EEs across all orders, including and up to the 6^th^ order (**Fig 4b**). To probe the expected number of EEs that should be successfully captured by RBA at the observed measurement error rate, we simulated 100 combinatorial landscape datasets with encoded experimental error ranging from SD = [0.01, 0.1] and log_10_ SMEs and EEs ranging from [0.1, 2.0]. Each dataset was simulated 1000 times, data were consolidated, and the resulting genotype-phenotype maps were analyzed using RBA (see **Methods**). We found that between 61–100% of SMEs and EEs were correctly classified as true positives across all orders, with a standard deviation of 0–10% across replicate simulations (**Fig. S2** and **S3**). False positives, at most, constituted less than 0.01% of the incorrectly assessed SMEs and EEs (**Fig. S4**), with the majority of errors being false negatives ranging from 0–39% (**Fig. S5**). The results of these simulations indicate that our experimental data, subjected to the error propagation pipeline, are highly robust to inaccuracies in assessing SME and EE significance.

The distribution of SMEs and EEs varied across substrates, and we found that variation within substrate classes was more pronounced than phenotypic data (**Fig. 4c**). Interestingly, we were able to identify a significant SME and EE for every mutation or interaction for at least one substrate (**Fig. 4c**). Previously observed trends were recapitulated, such as the positive effect of *f306***I** in the wt background solely for 2NH (0000x0 in **Fig. 4c**), or the positive effect of *d233***E**/*h254***R** for 2NH and acetate but not butyrate (xx0000 in **Fig. 4c**). Likewise, the strong and varied effect of *h254***R** seen in the PCA (**Fig. 2d**) and RFA (**Fig. 3**) was recapitulated in the breadth of EEs with *h254***R** across all substrates (**Fig. 4c-d**). We also computed the cross-correlations for each substrate pair between their SMEs and EEs and found that many of the correlations differed from those of the phenotypic changes (**Fig. 4e**). Correlations of 2NH with the other esters were much weaker (r = 0.33 for acetate and r = 0.26 for butyrate), while correlations across all OPs show more similarity, with SMEs and EEs of EPO appearing more correlated with the other OPs than the phenotypic changes (**Fig. 4e**). The strongest anticorrelation in the SMEs and EEs was between 2NH and EPO, likely explained by the iterative making and breaking of networks for each substrate, respectively, during the directed evolution for 2NH activity (**Fig. 4e**). We also observed greater correlations between the *p*-nitrophenyl substituted esters and the methyl-substituted OPs, particularly butyrate and MPT, at an *r* = 0.72, approaching the correlations seen in the within-OP class correlations (*r* = 0.74 of MPT and EPT). This suggests that although key effects may cause a strong divergence in observed activity of these two substrate classes, the network formation is similar, leading to effects such as diminishing losses in OPs and diminishing returns in the *p*-nitrophenyl esters. This is most apparent between DHC and acetate, where the phenotypic correlation (r = −0.60) differs greatly from the SME and EE correlation (r = 0.51), indicative of strong phenotypic divergence caused by a small subset of SMEs and EEs which, on average, are generally similar across the two substrates.

### Reference-based analysis enables dissection of intramolecular networks that underpin substrate trade-offs

In order to dissect key differences in intramolecular wiring during adaptation for 2NH, we probed two higher-order networks between three mutations for the esters and OPs: *d233***E**/*h254***R**/*l271***F** and *l271***F**/*f306***I**/*i313***F**. The *d233***E**/*h254***R** has been extensively studied for 2NH (Campbell et al., 2016; Tokuriki et al., 2012). Acetate showed a similar pattern of wiring to 2NH, with a strong SME from *h254***R**, a weaker *d233***E** SME, and a strong pairwise interaction for *d233***E**/*h254***R** (**Fig. 5**). Interestingly, this was not observed for butyrate, where the *h254***R** SME is 4–6x greater than the other esters, but is not supported by epistasis with *d233***E**; in fact, it exhibits 1.52-fold negative epistasis, suggesting the “bent” conformation is not preferred for *p*-nitrophenyl butyrate, but is for *p*-nitrophenyl acetate and 2NH (Buda et al., 2023; Campbell et al., 2016) (**Fig. 5**). Furthermore, the low idiosyncrasy of *l271***F** in butyrate may be explained by alternating signs of SMEs and EEs, where its initial tolerance is disrupted by negative epistasis, then again restabilized at higher-orders; a similar but opposite pattern is true for 2NH, while for acetate *l271***F** forms only one significant interaction out of the three that are present in the network (**Fig. 5**). This same three-way network also distinguishes between the four OPs. Firstly, as mentioned previously, *h254***R** is preferential for methyl-substituted OPs. This has been observed previously, and is likely the reason for this naturally-occurring polymorphism in the phosphotriesterase from *Agrobacterium radiobacter* (Jackson et al., 2005). In all cases, the pairwise effect of *d233***E**/*h254***R** is positive for all OPs, but it constitutes diminishing losses rather than an increase in activity, as it is always overshadowed by the negative contribution of *d233***E** (**Fig. 5**). Here, *l271***F** plays a larger role than in esters, with an extremely negative SME for EPT. We speculate that the larger ethyl- and thio-substituents are likely more sterically incompatible with the introduction of the bulkier **F***271*, though this seems to be strongly alleviated by *h254***R** for EPT, possibly stemming from the hydrogen-bond network formation between 254 and 271 as shown previously (Campbell et al., 2016). This alleviation is subtle in MPT, and actually negative for the oxo- substituted OPs, demonstrating a difference in active site shape preference between the substrates. The alleviation of *l271***F** in only the thio-substituted OPs is also mildly achieved by *d233***E**/*l271***F**; the mechanism for this is unclear due to their large distance with no opportunity to physically interact, thus implicating independent but synergistic remodeling of the active site better suited for thio-OPs, or an interaction between the mutations “relayed” through the substrate itself. This effect is flipped for the three-way network, with positive effects for the oxo-OPs and negative effects in the thio-OPs (**Fig. 5**). Thus, *d233***E**/*h254***R**/*l271***F** demonstrate network-based discrimination of thio- and oxo-substituted OPs.

**Figure 5.**
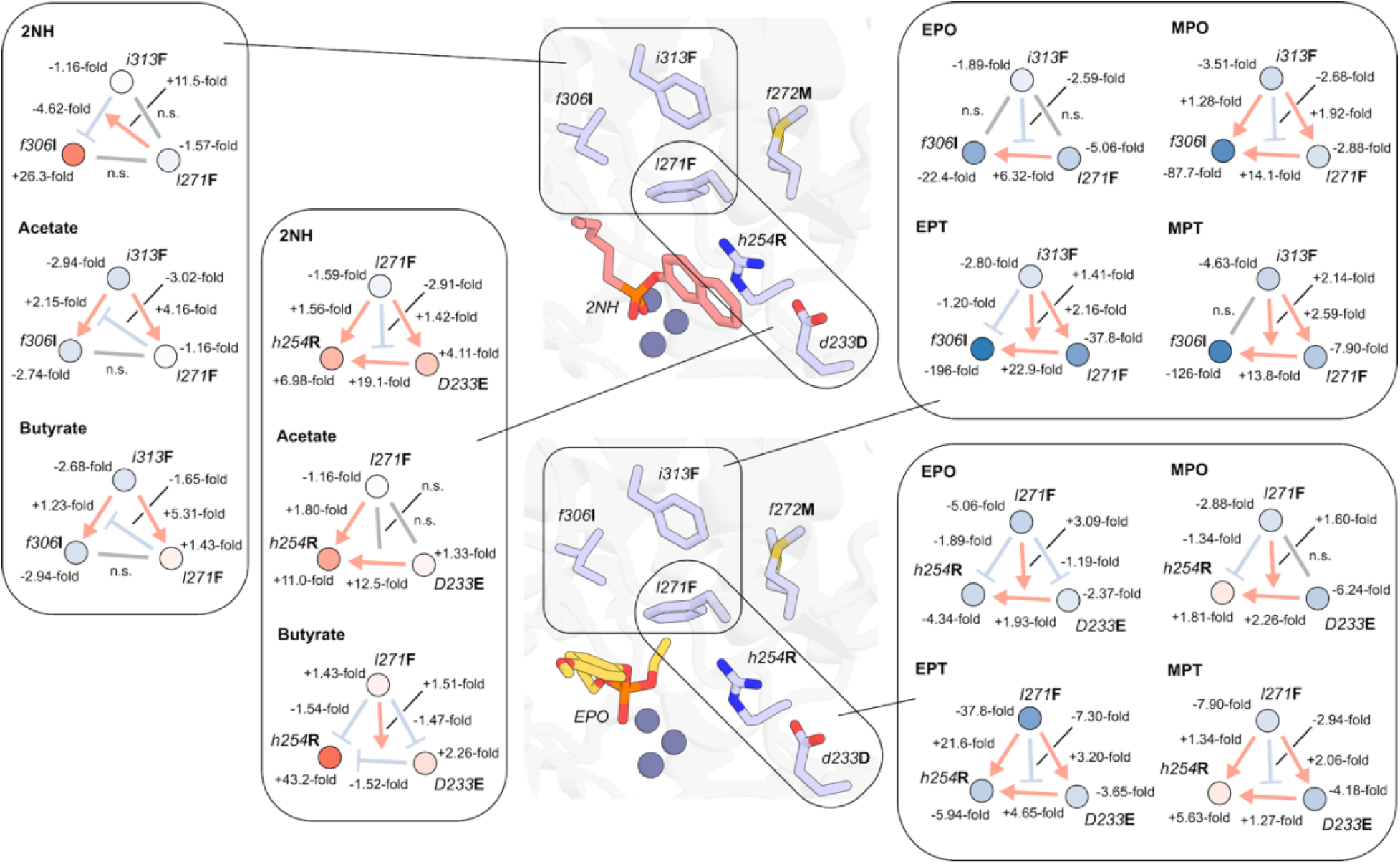
Higher-order intramolecular networks discriminate between different esters and organophosphates. The active site structure of phosphotriesterase (PTE) with the 2-naphthylhexanoate transition state analog (PDB: 4E3T; upper center) also overlayed with the ethyl-paraoxon transition state analog in the active site of *Agrobacterium radiobacter* PTE (PDB: 2R1N). Three-way networks of interest are highlighted, with insets representing an abstracted graph view of the network; nodes represent mutations and arrows indicate interactions (arrows pointing to arrows represent higher-order interactions), with red representing increases in activity, blue representing decreases, and grey representing no significant interactions.

For *l271***F**/*f306***I**/*i313***F**, we found the opposite trend for esters: acetate and butyrate showed similar wiring, distinct from 2NH. This is likely due to the unique effect of *f306***I** in 2NH. The positive SME *f306***I** is then disrupted by the negative EE of *f306***I**/*i313***F**, and finally restabilized by *l271***F** with no other pairwise EE contribution (**Fig. 5**). The presence of all three mutations appears to optimize the pocket proximal to the hexanoyl moiety of 2NH. This is not apparent for the shorter chain esters, with negative SMEs for *f306***I** and *i313***F**, and slight diminishing losses pairwise *f306***I/***i313***F** EE (**Fig. 5**). In contrast to 2NH, *l271***F/***i313***F** are positively epistatic and actually increase activity of acetate and butyrate relative to the wt background, and they do not tolerate the *l271***F/** *f306***I***/i313***F** network (**Fig. 5**). This may be indicative of beneficial electron density from three phenylalanine residues in this acyl chain pocket, which is better suited for the shorter chain esters. Despite the *p*-nitrophenyl substitution for acetate and butyrate, this may suggest that the substrate orientation of the two esters during catalysis is similar to 2NH, at least with respect to the acyl tail. As for the OPs, the network is best explained by the SME of *f306***I**, likely reducing stabilization of the *p*-nitrophenyl leaving group. Mutations at *l271***F** and *i313***F** compensate slightly, but not sufficiently to reverse the effect of *f306***I** (**Fig. 5**). One subtle difference in the network is revealed by the positive three-way effect of *l271***F***/f306***I/***i313***F** for thio- but not oxo- OPs (**Fig. 5**), again likely attributed to different physiochemical properties of the sulfur atom. Thus, the examination of these higher-order EEs reveals the gradual changes to the wiring of these active site intramolecular networks, which in turn dictates the recognition of different substrates and underpins the observed functional trade-offs.

## Discussion

This study demonstrates that the wiring of an enzyme’s active site is fundamentally multi-dimensional, relying on substrate-dependent interaction networks that can be precisely mapped through comprehensive epistatic analysis. By profiling the combinatorial landscape of six key mutations in PTE against nine diverse substrates, we have uncovered how intramolecular networks rewiring during adaptive evolution underpins activity trade-offs. This approach moves beyond global trends and focuses on revealing the idiosyncratic interactions that dictate substrate promiscuity.

The application of RFA across different substrates revealed that non-linear transformations necessitated the use of different coefficients for the sigmoidal fit for each substrate. Under the assumption that these transformations capture the non-linear mapping between mutational effects on intrinsic biophysical properties such as stability or expression, these coefficients should, in principle, remain identical regardless of the substrate being assayed, as they are a reflection of the free enzyme state. If this assumption is untrue, the present non-specific epistasis must be rooted in the differences in energies of the sub-states within the catalytic ensemble, which cannot be effectively captured by a non-linear transformation (Buda & Tokuriki, 2026). Indeed, previous work has demonstrated how differences in these sub-states can be revealed by varying ligand concentrations (Morrison et al., 2021; Morrison & Harms, 2022); it stands to reason that the same principle applies to varying substrate classes and types. However, these effects are deeply entangled with specific epistasis, making their isolation a significant challenge.

The implementation of RBA with rigorous error propagation, supported by extensive replicates, enabled a robust assessment of epistasis significance. Simulations suggest that even data with higher error rates remains amenable to RBA, as the primary source of error is the occurrence of false negatives rather than false positives, thus retained SMEs and EEs are likely true positives while discarded ones are more likely to be false negatives. Consequently, epistatic effects deemed significant through this pipeline are highly likely to represent genuine biological interactions, providing a reliable framework for identifying critical nodes and interactions in enzyme intramolecular networks. This statistical robustness is essential for distinguishing true higher-order interactions from the experimental noise often encountered in combinatorial landscapes.

Perhaps the most striking finding is the degree to which intramolecular wiring varies between substrates within the same chemical class, even when phenotypic adaptation appears similar. The divergence in epistatic patterns between the target substrate 2NH and its shorter-chain analogs, acetate and butyrate, highlights the network’s sensitivity to subtle changes in substrate moieties. For instance, the *d233***E**/*h254***R** pairwise interaction, the key driver of 2NH activity, is negative for butyrate. At face value, this implies that facilitating the “bent” active site conformation of **R***254*, preferred for 2NH, is suboptimal for shorter acyl chain esters or *p*-nitrophenyl substituted esters. However, the preservation of this positive effect in the *p*-nitrophenyl substituted acetate paints butyrate as a unique, idiosyncratic, outlier. Furthermore, the analysis of the *l271***F/***f306***I***/i313***F** network reveals a specialized pocket proximal to the hexanoyl moiety of 2NH. The unique positive synergy of these three residues for 2NH, contrasted with their varied effects on other esters and substrates, suggests that higher-order epistasis can reveal fine-tuning of the active site to accommodate specific features and moieties for the adaptive substrate.

These subtle, substrate-dependent differences in rewiring revealed by the RBA account for the emergence of trade-offs during evolution. This is best exemplified by the strong anti-correlation between both the epistatic effects as well as the phenotypic effects for 2NH and the native substrate EPO, which can be reconciled by the fact that accumulation of mutations to enhance hydrolysis of a dissimilar substrate is highly likely to disrupt the native catalytic network. However, this is not always the case. These trade-offs appear weaker when dissimilar substrates share some key similarities; methyl-substituted OPs and *p*-nitrophenyl esters, despite harboring different electrophiles, showed positive correlations in their epistatic effects, likely due to similar positioning within the active site as a result of the *p*-nitrophenyl moiety. Furthermore, despite the relatively weak correlation in the SMEs and EEs between 2NH and butyrate (**Fig. 4e**), their phenotypic correlation across the landscape were still positive (**Fig. 2b**), indicative of the esters’ tolerance for all six mutations. This suggests that despite the network’s discrimination between 2NH and butyrate, as exemplified by differences in EEs, phenotypic trade-offs can often be disproportionally controlled by a small subset of strong mutational effects of interactions (Muñiz-Trejo et al., 2025; Sailer & Harms, 2017). Thus, the use of intramolecular network analysis can help determine how rewiring can discriminate against, prioritize, or even tolerate certain substrate features during evolution, but discrepancies in network wiring between substrates should not be equated with significant differences in activity profiles.

In conclusion, substrate-dependent epistasis provides a powerful lens through which the subtle remodeling of enzyme active sites can be observed. This analysis enables grounded speculation on how rewiring events are underpinned by specific enzyme-substrate structural interactions. To verify these insights, future studies incorporating dynamic structural data or high-throughput MD will be essential to fully resolve the physical basis of the interactions identified here. We encourage the protein evolution community to adopt this error propagation method in order distill the key, relevant interactions, therefore streamlining the experimental workload required in resolving structures of mutational variants, and ultimately helping decode the complex intramolecular rewiring during evolution.

## Materials and Methods

### Combinatorial landscape measurements

During the directed evolution of phosphotriesterase toward arylesterase activity, we previously identified a cluster of six function-altering mutations and constructed all 64 (2^6^) genotypes (Miton et al., 2020). Previously, these combinations were tested for activity against *p*-nitrophenyl butyrate (Miton et al., 2020) and 2NH (Buda et al., 2023). Here, all variants were tested for activity against *p*-nitrophenyl acetate, ethyl-paraoxon (EPO), methyl-paraoxon (MPO), ethyl-parathion (EPT), methyl-parathion (MPT), dihydrocoumarin (DHC), and thiobutyl-butyrlolactone (TBBL). The 64 variants were constructed by site-directed mutagenesis and subcloned into a pET-27-Strep vector. Acyl phosphatase, cloned into the same vector, served as a negative control for all catalytic reactions. All variants were transformed into *E. coli* BL21(DE3) (Lucigen) carrying the pGro7 plasmid (Takara, Shiga, Japan) for GroEL/ES chaperones co-expression (Wyganowski et al., 2013). Variants were individually inoculated into 96-deep well plates containing lysogeny broth (LB) media, 100 μg/mL ampicillin, and 34 μg/mL chloramphenicol, then grown overnight at 30°C, 1050 rpm. Subsequently, 25 μL of overnight cultures were transferred to deep well plates containing 425 μL LB per well with 100 μg/mL ampicillin, 34 μg/mL chloramphenicol, 200 μM ZnCl_2_ and 0.2 % (w/v) of arabinose for chaperone co-expression. Cells were grown for ∼2 hrs at 37 °C until the OD reached ∼0.6. Expression was induced with 1 mM IPTG and cultures were incubated for 2 hours at 30°C. Cells were spun down at 4°C at 3,320 x g for 10 minutes and the supernatant was drained. Pellets were frozen for > 1 hr at −80 °C then resuspended in 200 μL of 50 mM Tris-HCl, 100mM NaCl, pH 7.5 buffer supplemented with 0.1% (w/v) Triton X-100, 200 μM ZnCl_2_, 100 μg/mL lysozyme (VWR) and 1 μL of benzonase (Novagen, 25 U/μL) per 100 mL of buffer. After 30 minutes of lysis, cell debris were spun down at 4°C at 3,320 x g for 30 minutes. If required, the clarified lysate was diluted (20–8,000x) prior to the activity assay to detect the initial linear phase of the reaction. Reactions were performed in 96-well plates (Corning Costar) containing 100 μL per well for > 1 hr. All substrates were dissolved in dimethyl sulfoxide, except TBBL, which was dissolved in acetonitrile. All substrates were freshly diluted in the lysis buffer prior to the assay, except DHC, which was diluted in 20 mM HEPES, pH 7.5. For acetate (Millipore Sigma) and butyrate (Millipore Sigma), initial rates were measured at 450 μM of substrate (at 405 nm). 2NH (Nanjing Sinfoo Technology Co.) hydrolysis was measured at 180 μM substrate, supplemented with 900 μM Fast Red TR salt (Sisco research laboratories); hydrolysis was monitored at 500 nm *via* complex formation with Fast Red. For all OPs (POE, POM, PTE, PTM; Millipore Sigma), initial rates were measured at 200 μM of substrate (at 405 nm). For DHC (Millipore Sigma), initial rates were measured at 360 μM of substrate (at 270 nm). For TBBL (a gift from the Tawfik Lab at the Weizmann Institute of Science), initial rates of hydrolysis were monitored at 180 μM of substrate, supplemented with 450 µM of Ellman’s reagent (DTNB; Millipore Sigma) (at 412 nm).

### Data processing pipeline

Initial rate replicates were grouped by genotype and normalized by (*i*) the mean wt initial rate for the respective substrate or (*ii*) the mean wt initial rate for EPO, then log_10_ transformed. Standard deviation of the mean and replicate number was collected for each genotype and substrate pair to compute the standard error of the mean. Relative mean absolute error (RMAE) was calculated by computing the mean of the sum of absolute errors for each genotype-substrate pair and dividing it the mean activity. The coefficient of variation (CV) was calculated by dividing the standard deviation by the mean activity. Raw data, code, and processed data have been deposited, available on GitHub (https://github.com/karolbuda/rba-error-propagation).

### Reference-free analysis

RFA was performed using the software provided by Park *et al*. (Park et al., 2024). We opted for the RFA-binary-fast functions without regularization to compare untransformed and sigmoidal transformations for increasing model orders up to 3^rd^ order. The Pearson coefficients were grouped for each order across all substrates, and a paired t-test was performed to determine the differences in fit. For *T*_90_ assessment, we used a 2^nd^ model order for each substrate.

### Reference-based analysis

SMEs and EEs were computed as previously described (Buda et al., 2023). Briefly, genotypes were represented as binary strings *_g_* ∈ {0,1}*^N^*. Let *_f_*(*_g_*) represent the normalized, log_10_-transformed function. We defined the SME at position *_i_* as the first-order difference:

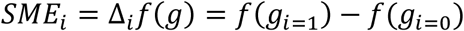

Where *_gi_*_=1_ and *_gi_*_=0_represent the derived and ancestral states at position *i* within a constant genetic background, respectively. Pairwise EEs were then calculated as successive difference operators:

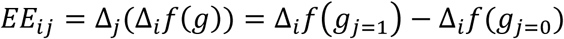

Generally, an *n*^th^ order interaction among a set of positions *S* is given by:

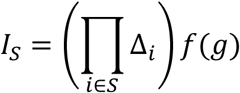

Then, to compute the significance of each SME and EE, the standard error of the mean was propagated for each constituent *_f_*(*_g_*) and computed the effective sample size of the SME or EE by calculating the harmonic mean of the replicates for each constituent *_f_*(*_g_*). For each SME and EE, a t-value was computed, as well as an adjusted p-value using the false discovery rate method for the number of SMEs and EEs tested.

### Simulations for reference-based analysis assessment

A single substrate dataset of log_10_ activities was simulated for each genotype based on predefined SMEs and EEs. We initially defined the values of SMEs = 1 and EEs = 0 and simulated phenotypes for all combinatorial genotypes across six positions. For each genotype, we randomly sampled 3 points from a gaussian distribution with mean equal to the “true” activity and standard deviation varied from [0.01, 0.02, 0.03, 0.04, 0.05, 0.06, 0.07, 0.08, 0.09, 0.1]. We ran the simulations defining SMEs and EEs at each order as [0.1, 0.2, 0.3, 0.5, 0.7, 1.0, 1.3, 1.5, 1.7, and 2.0], keeping SMEs = 1 and other EEs = 0, where relevant (*e.g.*, at order 2, SMEs were all 1, pairwise EEs were varied, and all higher-order EEs were 0). We repeated all simulations 1000 times and consolidated the data. We computed the mean true positives, false positives, and false negatives for all SMEs and EEs as a percentage from the 1000 simulations, also recording the percent standard deviation across all simulations. Simulation code and data were deposited in GitHub (https://github.com/karolbuda/rba-error-propagation).

## Supplementary material description

Supplementary file 1: Supplementary figures for non-linear transformation and reference-based analysis simulations

## Acknowledgements

K.B. and N.T. are supported by the Natural Sciences and Engineering Research Council of Canada (NSERC; https://www.nserc-crsng.gc.ca/index_eng.asp)/Discovery Grants Program (RGPIN-2023-05135). The funders had no role in study design, data collection and analysis, decision to publish, or preparation of the manuscript. We also thank Dr. Miriam Kaltenbach for the technical assistance with variant construction. Finally, we thank all of the Tokuriki lab members for valuable discussions.

