## Supplementary figures for "Substrate-dependent epistasis probes active site intramolecular wiring"

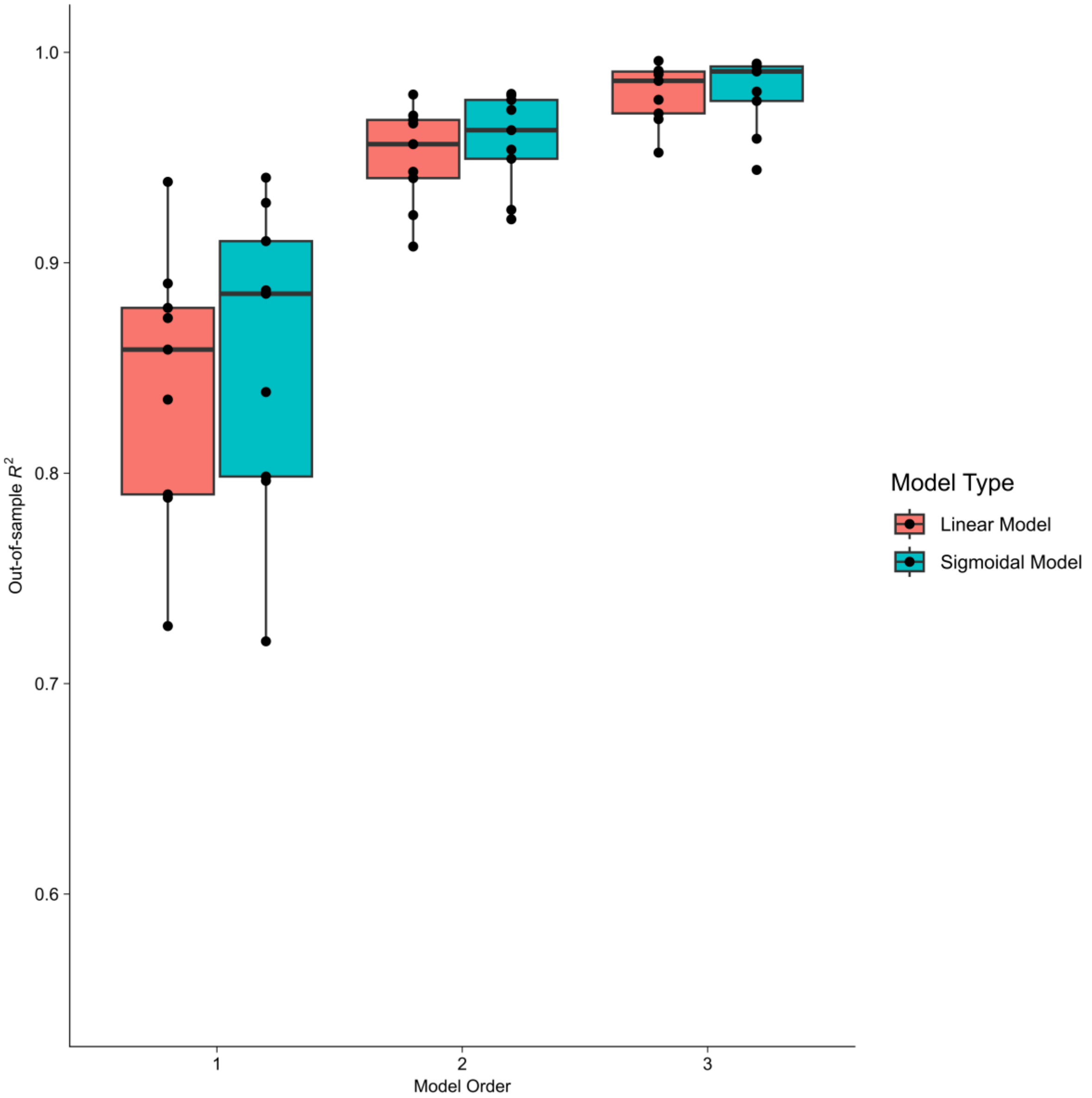


**Figure S1 – Negligible model improvement in reference-free analysis fit from sigmoidal non-linear transformation.** Out-of-sample $R^2$ predicted from coefficients inferred from an 80-20% train-test split. Linear model used no transformation of phenotype to genetic score; sigmoidal model used a two-parameter fit outlined in Park *et al.* (2024).

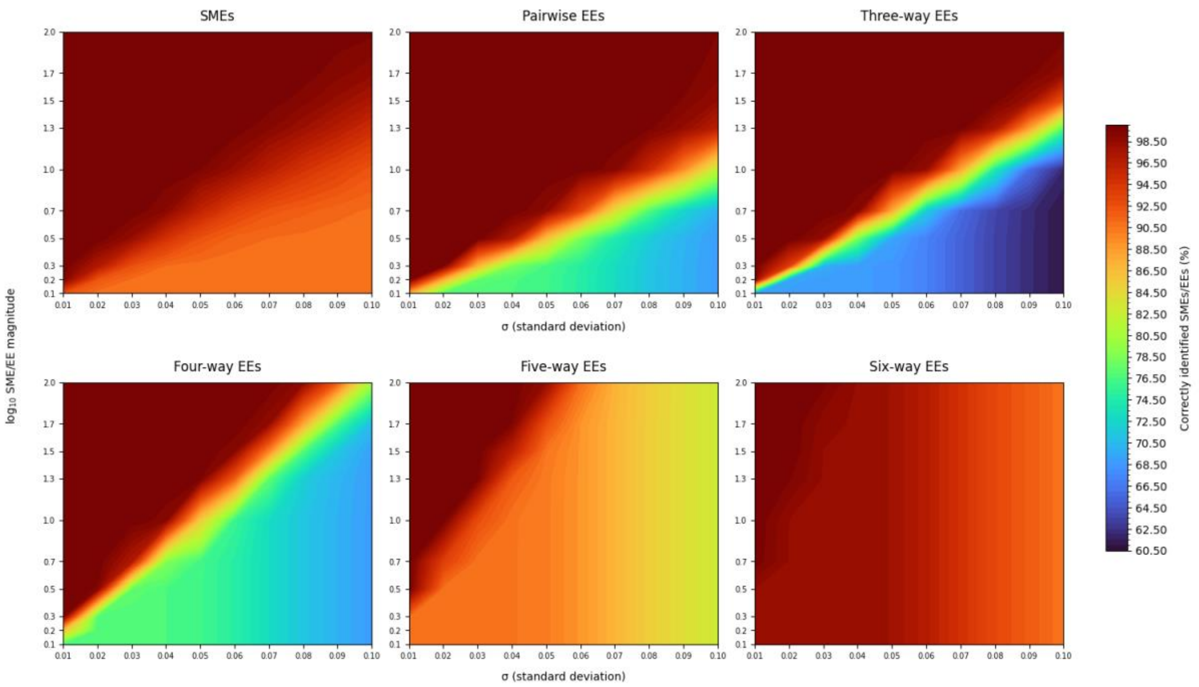


**Figure S2 – Mean true positive % of SMEs and EEs as detected by reference-based analysis with error propagation on simulated data.** Simulated errors were represented as standard deviation (x-axis) and SMEs and EEs were uniformly assigned at order (y-axis). Replicates were performed 1000 times.

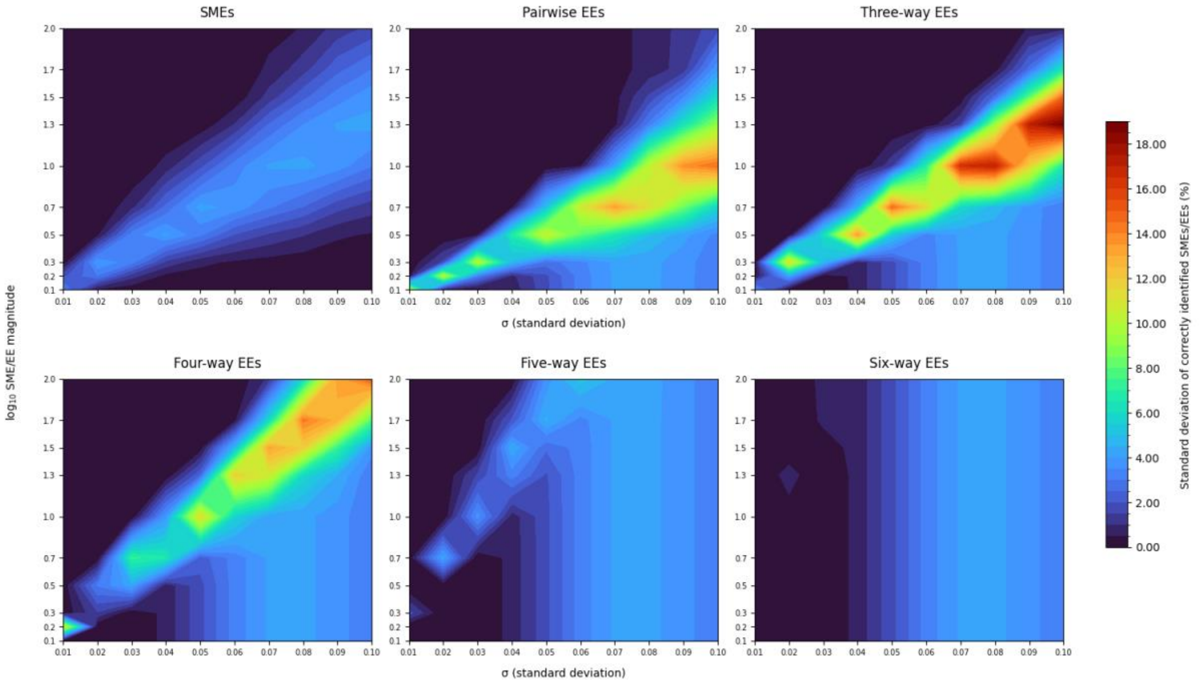


**Figure S3 – Standard deviation of true positive % of SMEs and EEs as detected by reference-based analysis with error propagation on simulated data.** Simulated errors were represented as standard deviation (x-axis) and SMEs and EEs were uniformly assigned at order (y-axis). Replicates were performed 1000 times.

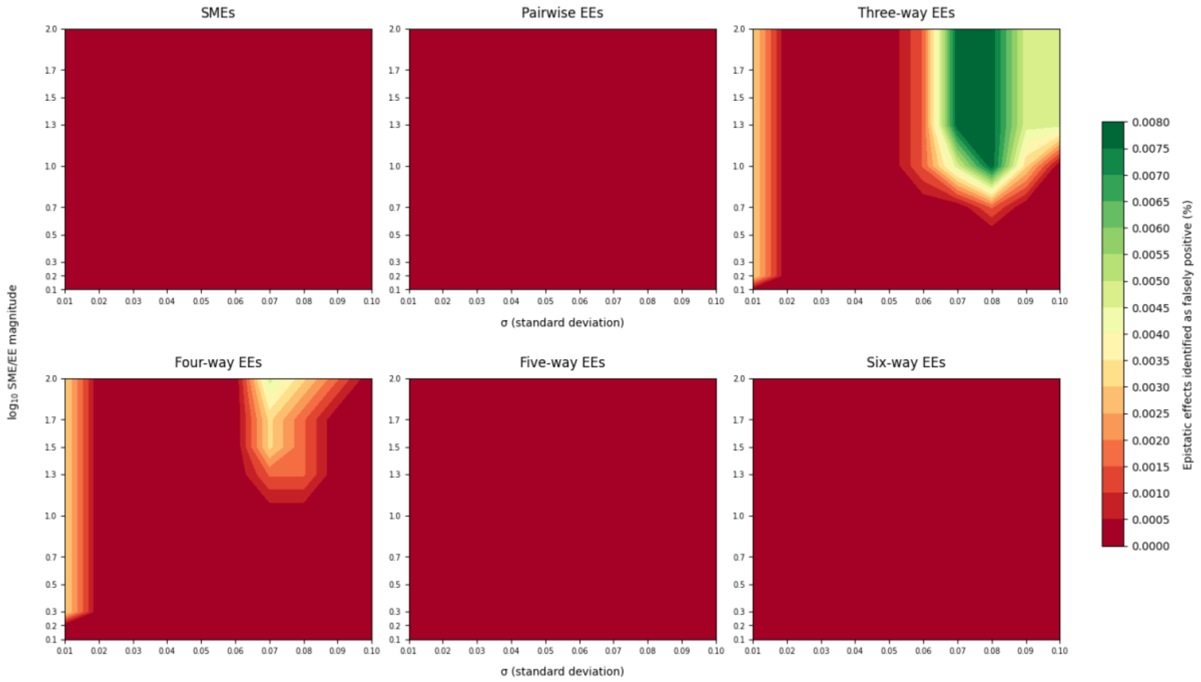


**Figure S4 – Mean of false positive % of SMEs and EEs as detected by reference-based analysis with error propagation on simulated data.** Simulated errors were represented as standard deviation (x-axis) and SMEs and EEs were uniformly assigned at order (y-axis). Replicates were performed 1000 times.

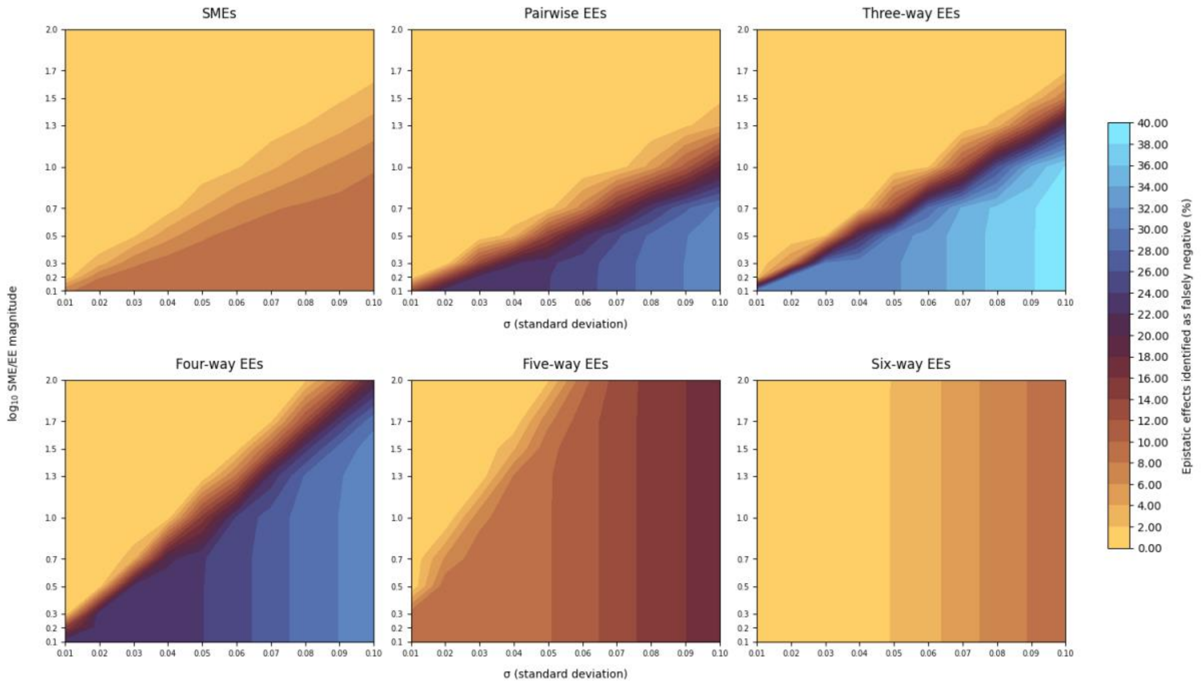


**Figure S5 – Mean of false negatives % of SMEs and EEs as detected by reference-based analysis with error propagation on simulated data.** Simulated errors were represented as standard deviation (x-axis) and SMEs and EEs were uniformly assigned at order (y-axis). Replicates were performed 1000 times.
